# Examining the effect of chronic intranasal oxytocin administration on the neuroanatomy and behavior of three autism-related mouse models

**DOI:** 10.1101/2021.01.13.426562

**Authors:** Zsuzsa Lindenmaier, Jacob Ellegood, Monique Stuive, Kaitlyn Easson, Yohan Yee, Darren Fernandes, Jane Foster, Evdokia Anagnostou, Jason P. Lerch

## Abstract

Although initially showing great potential, oxytocin treatment has encountered a translational hurdle in its promise of treating the social deficits of autism. Some debate surrounds the ability of oxytocin to successfully enter the brain, and therefore modify neuroanatomy. Moreover, given the heterogeneous nature of autism, treatment will only amerliorate symptoms in a subset of patients. Therefore, to determine whether oxytocin changes brain circuitry, and whether it does so variably, depending on genotype, we implemented a large randomized, blinded, placebo-controlled, preclinical study on chronic intranasal oxytocin treatment in three different mouse models related to autism with a focus on using neuroanatomical phenotypes to assess and subset treatment response. Intranasal oxytocin (0.6IU) was administered daily, for 28 days, starting at 5 weeks of age to the *16p11*.*2* deletion, *Shank3* (exon 4-9) knockout, and *Fmr1* knockout mouse models. Given the sensitivity of structural magnetic resonance imaging (MRI) to the neurological effects of interventions like drugs, along with many other advantages, the mice underwent *in vivo* longitudinal and high-resolution *ex vivo* imaging with MRI. The scans included three *in vivo* T1-weighted, 90*u*m isotropic resolution scans and a T2-weighted, 3D fast spin echo with 40*u*m isotropic resolution *ex vivo* scan to assess the changes in neuroanatomy using established automated image registration and deformation based morphometry approaches in response to oxytocin treatment. The behavior of the mice was assessed in multiple domains, including social behaviours and repetitive behaviours, among others. Treatment effect on the neuroanatomy did not reach significance, although the pattern of trending effects was promising. No significant effect of treatment was found on social behavior in any of the strains, although a significant effect of treatment was found in the *Fmr1* mouse, with treatment normalizing a grooming deficit. No other treatment effect on behavior was observed that survived multiple comparisons correction. Overall, chronic treatment with oxytocin had limited effects on the three mouse models related to autism, and no promising pattern of response susceptibility emerged.

## 2 Introduction

Autism spectrum disorder (ASD) is a very heterogeneous neurodevelopmental disorder, characterized by impaired social communication and repetitive behaviors. To date, there is no pharmacological treatment to improve the social/communication deficits in autism patients (Anagnostou et al. (2012); de la Torre-Ubieta et al. (2016)). One promising pharmacological treatment for these social communication deficits is oxytocin.

Oxytocin is a neuromodulator, used in humans commercially to induce labor, but is known for its social effects. Oxytocin has been shown to increase activity in brain structures involved in processing socially meaningful stimuli (Andari et al. (2020, 2016); Aoki et al. (2014); Aoki and Yamasue (2015); Benner et al. (2021); Domes et al. (2013); Gordon et al. (2013); Watanabe et al. (2014, 2015)). Oxytocin also has numerous brain targets, including but not limited to the amygdala, hippocampus, hypothalamus, and nucleus accumbens (Quintana et al. (2019)). Therefore, it may be postulated that treatment with oxytocin would normalize social behaviors and memory, while decreasing anxiety and parasympathetic activation (fear/stress) (Benarroch (2013); Guastella and Hickie (2016)).

Oxytocin treatment has recently also been considered in autism for reasons other than its social effects (Anagnostou et al. (2012); de la Torre-Ubieta et al. (2016); Guastella and Hickie (2016)), including altered quantities of markers of oxytocin (i.e. mRNA, blood levels, etc) in individuals with autism when compared to typically developing controls(Andari et al. (2020); Bittel et al. (2006); Green et al. (2001); Gregory et al. (2009); Hernandez et al. (2017); Jacob et al. (2007); Modahl et al. (1998); Sebat et al. (2007)).

A number of existing studies have investigated the effects of oxytocin in autism and related animal models, with varied results. Many acute studies of oxytocin showed promise, normalizing social deficits and/or repetitive behaviours in mouse models of autism (Graustella and MacLeod (2012); Guastella and Hickie (2016); Huang et al. (2014); Peñagarikano et al. (2015); Sala et al. (2011); Teng et al. (2013)). One study in rodents found opposing results for acute vs chronic oxytocin administration (Huang et al. (2014)), while another specifically focusing on the social effects of oxytocin, found that chronic administration actually hinders some social behaviors in prairie voles (Bales et al. (2014)). Only three clinical trials found improvements with chronic administration, all with relatively low sample sizes (n=18, 19, 30)(Anagnostou et al. (2012); Watanabe et al. (2015); Yatawara et al. (2016)), while most recently, a large clinical trial (n=250) showed no significant effect of treatment on social or cognitive functioning over a period of 24 weeks (Sikich et al. (2021)). As Huang et al. (2014) put it, there seems to be a “translational hurdle” in the movement of oxytocin from acute studies to chronic. Despite the promise of acute treatment, given the lifelong nature of the disorder and its symptoms, chronic treatment is likely the only way amelioration of autism symptoms would occur.

Moreover, there is a gap in our knowledge surrounding oxytocin’s effects on the brain. A number of neuroimaging studies investigating oxytocin’s effect on neural activity during behavioral tasks found enhanced brain activity predictive of behavioral effects due to treatment (Andari et al. (2020, 2016); Aoki et al. (2014); Aoki and Yamasue (2015); Benner et al. (2021); Gordon et al. (2013); Watanabe et al. (2014, 2015)). But these effects are 1) from a diverse number of regions across the brain, including the amygdala, the anterior cingulate cortex, the anterior insula, the hippocampus, the medial prefrontal cortex, the striatum, and others, 2) from studies with low sample sizes (<40 participants) and 3) were observed using acute oxytocin administration, leaving our understanding of where the drug acts (and modulates downstream) after administration limited, despite knowing where oxytocin receptors are located (see Figure 2) (Benarroch (2013); Gould and Zingg (2003); Quintana et al. (2019)) and our understanding of some of the effects oxytocin has on neural activity in the context of social behavior.

This problem is further exacerbated by the pharmacology of oxytocin. The neuropeptide is too large to cross the blood brain barrier and has a very short halflife (Neumann et al. (2013)). Therefore, an administration method is needed that quickly and directly allows oxytocin to enter the brain. Though methods like intracerebroventricular (ICV) infusion and peripheral injections have been used in animal models, there is a need for a method that is easily employed by humans (as opposed to the intravenous (IV) lines used for labour induction). Intranasal administration allows for simple, non-invasive treatment that circumvents the issues with the blood brain barrier and is easily used both in humans and in mice (Neumann et al. (2013)). Although recent literature suggests this is a promising mode of administration (Galbusera et al. (2017); Pagani et al. (2020)), some debate surrounds whether oxytocin really crosses the blood brain barrier. For example, Leng and Ludwig (2016) argued that the treatment effects observed in studies assessing intranasal oxytocin (like in Neumann et al. (2013)) were solely due to peripheral treatment effects. A method that accurately assesses oxytocin’s ability to enter the brain is needed to settle this debate.

Importantly, given the substantial heterogeneity of ASD, there is a need for studies to delineate phenotype-based subsets that may respond to oxytocin treatment (Ecker and Murphy (2014); Guastella and Hickie (2016)). This is perhaps best demonstrated by Engelman and colleagues (2008) in their co-clinical study of lung cancer (Engelman et al. (2008)). A clinical trial of a promising therapeutic failed to show treatment effect in their population. They simultaneously treated three different genetic knockout mouse strains, with the same treatment used in humans. The group realized that two of the strains responded 100% to the treatment, while one did not. Armed with this genetic information, they divided the clinical population into subsets by genotype and found that, in fact, all the patients not carrying the genetic mutation of the non-responder mouse model improved with the treatment. To this end, a study is necessary that incorporates numerous phenotypes across multiple mouse lines, and also makes use of multivariate analyses to inform clinical trials, all in the context of ASD.

Numerous studies have spoken to the utility of magnetic resonance imaging (MRI) in the stratification of ASD individuals into biologically homogeneous subgroups (Bernhardt et al. (2016); Ecker and Murphy (2014); Hong et al. (2018); Hrdlicka et al. (2005); Kushki et al. (2019)). Magnetic resonance imaging (MRI) offers the ability to quantify the structural phenotype of both mice and humans, allowing for whole brain coverage and translatability to human imaging studies, both in acquisition and analysis (Chen and Nieman (2011); Hong et al. (2018); Lerch et al. (2011); Nieman et al. (2018); Szulc et al. (2015)). It also affords the additional advantages of high-throughput, having a link with behavior (Anacker et al. (2016); Ellegood et al. (2015, 2013); Lerch et al. (2011); Nieman et al. (2007)), and showing evidence of plasticity with exposure (Vousden (2018); Zhang et al. (2018)).

Perhaps most importantly, MRI has shown sensitivity to the neurological effects of interventions like drugs (Bade et al. (2017); Dudek et al. (2015, 2016); Lu et al. (2007); Perrine et al. (2015); Scholz et al. (2015); van der Plas et al. (2020); Vousden (2018); Wheeler et al. (2013); Zhang et al. (2018)). These studies point to the utility of using MRI to characterize structural plasticity in response to pharmacology. This allows the ability to assess whether oxytocin modifies brain circuits and whether it does so variably depending on genotype.

To explore if we could understand patterns of response in mice, we executed a mouse study in parallel, akin to a co-clinical trial. Specifically, a large randomized, blinded, placebo-controlled, preclinical study on chronic intranasal oxytocin treatment in three different mouse models related to autism was performed with a focus on using neuroanatomical phenotypes to assess and subset treatment response. A wide range of behavioural measures was also assayed in a high-throughput method of treatment evaluation.

## 3 Methods

### 3.1 Subjects

Three in-bred strains of autism mouse models were used: the *16p11*.*2* heterozygous knockout (B6129S-Del(7Slx1b-Sept1)4Aam/J; JAX #013128) (Horev et al. (2011))(henceforth referred to as “*16p*”), the *Fmr1* hemizygous knockout (FVB.129P2Pde6b+ Tyrc-ch Fmr1tm1Cgr/J; JAX #004624)(Bakker et al. (1994))(henceforth referred to as “*Fmr1* “), and the *Shank3* exon 4-9 (ANK domain) homozygous knockout (B6(Cg)-Shank3tm1.2Bux/J; JAX #017890)(Bozdagi et al. (2010))(henceforth referred to as “*Shank3* “). The mice were on the following background strains, respectively: B6129SF1/J (JAX #101043)(henceforth referred to as “129”), FVB.129P2-Pde6b+ Tyrc-ch/AntJ (JAX #004828)(henceforth referred to as “FVB”), C57BL/6J (JAX #000664)(henceforth referred to as “C57”).

These strains were chosen because they were genetic models with high face validity. Specifically, they are models with strong analogies to the human condition. For example, the *Fmr1* mouse line is a well studied model of the most common single gene mutation related to autism. More pragmatic considerations like availability were also considered, along with the models’ distributions in the clusters from Ellegood et al. (2015).

Active colonies for each strain were maintained at The Centre for Phenogenomics (TCP). Breeders were set up to yield control litter-mates for each strain. Specifically, for the *16p* strain, heterozygous mice were bred with wildtype 129 mice to yield the desired heterozygous mutants and wildtype control litter-mates. For the *Fmr1* strain, heterozygous females were bred with wildtype males to yield hemizygous males and wildtype control litter-mates. For the *Shank3* mice, two heterozygous mice were bred to yield homozygous *Shank3* mutants and wildtype control littermates.

Mice were group housed, and an attempt was made to achieve a sample size of 14 male mice per genotype, per strain, per treatment (Table 2). The sample size was chosen based off a power analysis conducted *a priori* that determined that 14 subjects in each group should be sufficient to detect group differences, based off the variability in the chosen neuroanatomical phenotypes. Data was collected and analyzed for all mice for all behavioral tests, except for sociability. Although the software appeared to be recording, the authors later discovered that one of the laptops failed to acquire the sociability data, leading to a loss of approximately 40% of the social behavior data (reduced sample sizes are displayed in Table 2b).

All studies and procedures were approved by the TCP Animal Care Committee, in accordance with recommendations of the Canadian Council on Animal Care, the requirements under the Animals for Research Act, RSO1980, and the TCP Committee Policies and Guidelines. Mice were maintained under controlled conditions (25 C, 12 hour cycle) at TCP in sterile, individually ventilated cages and were provided standard chow (Harlan, Teklad Global) and sterile water *ad libitum* via an automated watering system.

### 3.2 Experimental Time-line

The experimental time-line is outlined in Figure 1. At five weeks of age, mice underwent *in vivo* manganese-enhanced magentic resonance imaging (MEMRI) scanning and were randomly assigned to a treatment group (oxytocin or placebo) such that the experimenter remained blinded. Mice were treated everyday, for 28 days total. After an *in vivo* MEMRI scan at postnatal week 8, mice were assessed on four different behavioral paradigms, while still receiving treatment. At nine weeks of age, the mice were scanned with *in vivo* MEMRI once more, perfused, and scanned *ex vivo* at a later time-point. The study was designed to be an exploratory study and therefore no primary outcome was defined before the study started.

**Figure 1:**
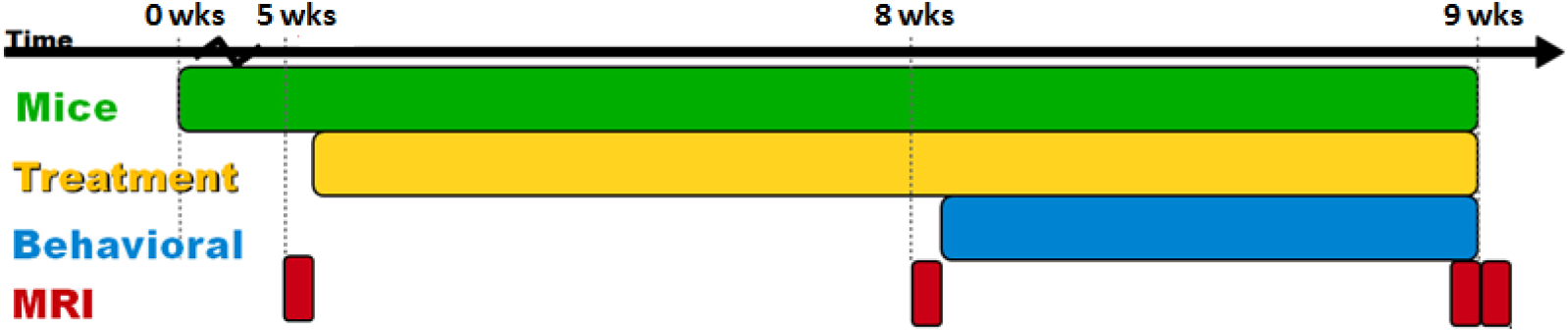
Time-line illustrating the project design. Green indicates the life of the mice, yellow indicates treatment administration, blue indicates period of behavioral testing, while red indicates occurrences of magnetic resonance imaging (MRI).

The protocol was designed to maximize throughput, and therefore the measures chosen were quick to perform, with little intervention or lasting effects. The behavioral tests were conducted in proximity to each other to minimize the variability between the brain and behavioral phenotypes (Crawley (2007); Paylor et al. (2006)). Similarly, the age of the mice was chosen so that single day variations would not have large effects on brain or behavioral phenotypes. The length of treatment was chosen to probe chronic treatment effects, while remaining manageable for highthroughput. To increase replicability, the *ex vivo* scan was performed to parallel the protocol from Ellegood et al. (2015). Great care was taken in the conception of this project to ensure that effects observed would be due to treatment and not confounding effects. Pilot testing involving wildtype mice was employed to ensure expected neuroanatomical and behavioral results before the main study commenced.

Although it is typical to space behavioral tests apart by several days, the methods employed were developed to increase throughput to maximize the number of subjects in the study, thereby increasing the statistical power to detect a treatment effect (Crawley (2007); Paylor et al. (2006)). Data from this study was compared to the literature as well as previous data acquired from the lab to ensure that the distribution of the data was expected. Specifically, if an effect of methodology was present, the distribution of the data from later behavioral tests (i.e. Open Field) would skew from the expected distribution based off previous literature (where the test was employed alone or spaced apart from other tests). Expected differences due to genotype (i.e. placebo-treated *16p* vs placebo-treated 129 control) would also be altered if an effect of protocol was present. Other measures were additionally implemented to ensure minimal interference; for example, the order the behavioral tests were performed was chosen based off studies investigating effects of training history (Crawley (2007); McIlwain et al. (2001)).

### 3.3 Oxytocin Treatment

Oxytocin (Sigma-Aldrich) was dissolved in saline and administered intranasally everyday for 28 days. Each morning, between 10am and 11am, the mice received 0.15*µ*g/10*µ*L. This dose was chosen to be 0.6IU, too low to produce peripheral effects and similar to the quantity used in humans (Bales et al. (2013); Meziane et al. (2015); Neumann et al. (2013)). Control mice received the same quantity of saline (placebo). A similar protocol was used as Huang et al (2014) (Huang et al. (2014)): a 200-ul Eppendorf pipette was used for administration, and drops of the solution were gently placed equally on both nostrils of each mouse. The primary experimenter was blinded to treatment and genotype, and was only unblinded at the time of analysis, when experiments were over.

### 3.4 Neuroanatomical Phenotype

Mice were injected in their intraperitoneal (I.P.) cavity with a manganese (Mn) contrast agent (0.4 mmol/kg dose of 30mM MnCl2 solution) up to 24 hours before the scan (Wadghiri et al. (2004)). A multi-channel 7.0T magnet (Agilent Technologies) was used that allows up to 7 animals to be scanned simultaneously. The mice were anesthetized using 1% isoflourane during T1-weighted scanning with 90 *u*m isotropic resolution (TR=29ms, TE=5.37ms, 1 hour 52 minute scan time). The animals’ breathing and temperature were monitored. Mice were imaged at 5, 8, and 9 weeks of age.

One *ex vivo* MRI scan was performed at least a month after the mice were sacrificed (Elizabeth De Guzman et al. (2016)). The procedure followed closely follows the procedure described in Lerch et al. (2011) and Nieman et al. (2018). To prepare the brains for scanning and enhance contrast, a fixation and perfusion procedure was done as previously described in Cahill et al. (2012). The same multichannel 7.0T magnet (Varian Inc, Palo Alto, CA) was used, with the ability to scan up to 16 brains simultaneously. A T2-weighted 3D fast spin echo sequence was used to yield an isotropic resolution of 40 *u*m (cylindrical acquisition, TR=350ms, TE=12ms, 14 hour scan time)(Spencer Noakes et al. (2017)).

Data from both imaging types was analyzed using image registration and deformation based morphometry approaches as described in Lerch et al. (2011). Briefly, images were linearly (6 parameter, then 12 parameter) and nonlinearly registered together. At completion of this registration, all scans were deformed into alignment with each other in an unbiased fashion. These registrations were performed with a combination of MNI autoreg tools (Collins et al. (1994)) and ANTS (advanced normalization tools)(Avants et al. (2008, 2011)). The changes within regions and across the brain can be examined using deformation based morphometry (DBM) and MAGeT (a multi-atlas registration-based segmentation tool)(Chakravarty et al. (2013)). An atlas that segments 159 structures in the adult mouse brain was used for the *in vivo* image analysis, and the *ex vivo* analysis has further delineations resulting in a total of 182 distinct regions (Dorr et al. (2008); Qiu et al. (2018); Richards et al. (2011); Steadman et al. (2014); Ullmann et al. (2013)). Segmentations of all brains passed quality control by visual inspection (see Table 1).

**Table 1:** Table depicting the number of images that passed quality control at each imaging time point in each strain, broken down by treatment. A minimum of n=14 was aimed for per genotype, per strain, per treatment. Although all mice were scanned at all time points, some images were excluded due to motion artifacts or low SNR.

| Timepoint | 5wks |  | 8wks |  | 9wks |  | *Ex vivo* |  |
| --- | --- | --- | --- | --- | --- | --- | --- | --- |
| Strain | Oxy | Pl | Oxy | Pl | Oxy | Pl | Oxy | Pl |
| *16p* | 6 | 6 | 6 | 9 | 10 | 10 | 15 | 15 |
| 129 | 3 | 9 | 7 | 8 | 9 | 12 | 14 | 15 |
| *Fmr1* | 10 | 8 | 11 | 9 | 11 | 7 | 15 | 14 |
| FVB | 14 | 14 | 14 | 12 | 12 | 10 | 16 | 13 |
| *Shank3* | 19 | 16 | 19 | 16 | 18 | 12 | 19 | 17 |
| C57 | 13 | 11 | 14 | 12 | 12 | 12 | 14 | 12 |

#### 3.4.1 Statistics

Data from both imaging methods was analyzed using the R statistical language (R Core Team (2017)) using linear mixed effects models (Lmer)(Bates et al. (2015); Pinheiro and Bates (2000)). Linear mixed effects models were used because they incorporate both fixed and random effects and are useful for longitudinal data, as well as unbalanced study designs (Bates et al. (2015); Pinheiro and Bates (2000))(see (Vousden et al. (2018)) and (Qiu et al. (2018))for fuller explanations on similar data).

In the case of the imaging data, linear mixed effects models were used to determine the effect of genotype (defined as wildtype, *16p, Fmr1*, or *Shank3*) and treatment on the volume for each region of interest (ROI) or voxel (Equation 1). Random intercepts for each background strain were incorporated. A log-likelihood test was performed to assess the significance of treatment and genotype, and their interaction. Data was analyzed as absolute volumes as well as relative volumes by covarying for total brain volume (Mankiw et al. (2017)).

To assess the effect of treatment on time, brain region volume at the 5 weeks *in vivo* scan point was used to predict, along with genotype and treatment, the latter brain region volumes (Equation 2). For simplicity, relative volumes were calculated as proportion of total brain volume. Additional information about the genotype effects was also analyzed by subsetting by background strain and assessing the main effects of genotype (mutant versus wildtype) and treatment using linear models in the *ex vivo* dataset (Equation 3). Multiple comparisons in the data from all analyses was corrected for using false discovery rate (FDR) correction (Genovese et al. (2002)), with p-values pooled by treatment, genotype, and interaction independently.

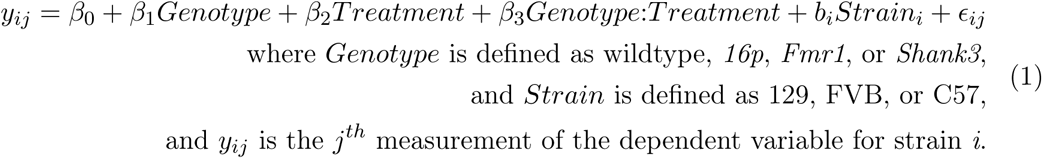

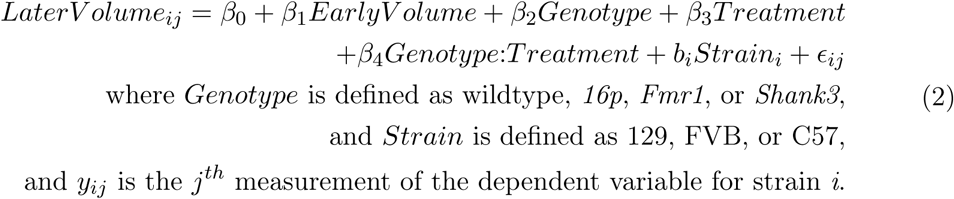

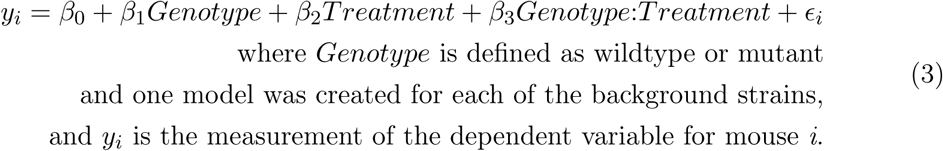

#### 3.4.2 Spatial gene expression analysis

To identify potential mechanisms that might cause these volume changes, we linked spatial gene expression data to the MRI-determined phenotypes. Under the assumption that if a gene is associated with volume change, then subsequent volume changes are spatially restricted to regions in which that gene is expressed, we sought to identify if OXTR gene expression correlated with volume changes due to oxytocin treatment.

We used the Allen Mouse Brain atlas from the Allen Institute, that profiled 4345 expression images in adult mice (P56)(limited number of genes due to the use of the coronal dataset only)(Lein et al. (2007)). Spatial correlation between structural volume and gene expression at each voxel were computed. The brain was subdivided into ten regions with increasing gene expression profiles. These subdivisions were ranked by correlation to the OXTR linear mixed-effects model (Equation 1) to assess which subdivision’s expression is most similar and most dissimilar to the observed volumetric changes.

These data processing and spatial enrichment methods have been previously used elsewhere, both in exploratory methods (Fernandes et al. (2017); **?**), and to support targeted hypotheses (Carriere et al. (2020)).

### 3.5 Behavioral Phenotype

On the third week of treatment, the mice were assessed for repetitive behaviors (as assessed by grooming), social behaviors (three chamber sociability task), anxiety and hyperactivity (open field), learning and motor coordination (rotarod), and memory (novel object recognition).

#### 3.5.1 Statistics

Behavioral data was analyzed using the R statistical language (R Core Team, 2016)(R Core Team (2017)). As with the imaging data, the behavioral data was assessed in two ways, to look at a general treatment effect across genotypes and background strains, as well as within strain. In this case, linear mixed effects models were created and compared using log-likelihood tests to determine the effect of genotype (defined as wildtype, *16p, Fmr1*, or *Shank3*) and treatment, and their interaction on each behavioral test (see Equation 4). Random intercepts for each background strain were incorporated.

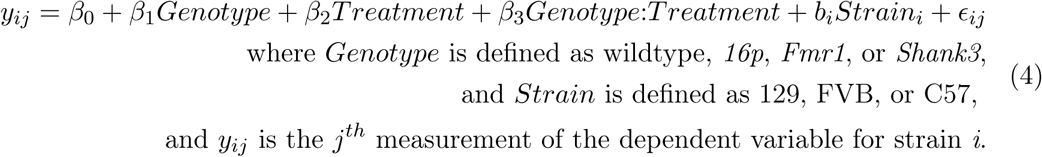

Additional information regarding treatment response and genotype effect was assessed by subsetting by background strain (129, FVB, C57) and using traditional linear regression analysis (see Equation 5). For each strain, separate linear models were created and compared using ANOVA, to determine whether a particular fixed effect (treatment, genotype, or their interaction) significantly i mproved t he model. The distribution within strain of each behavioral test was also assessed and corrected for, before analysis. The genotype effect was used to compare to existing behavioral literature. All data was corrected for multiple comparisons using false discovery rate (FDR) correction (Genovese et al. (2002)), with q-values pooled by treatment, genotype, and interaction independently.

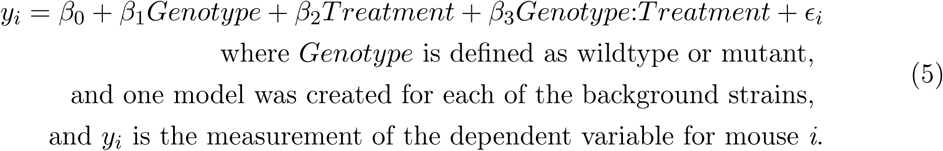

To explore the multivariate nature of the project, principle component analysis (PCA) was employed. Due to the reduced sample size in the sociability data, analyses were done on two sets of the data: 1) all mice that had grooming, open field, and rotarod data (“all mice, no sociability data”)(see Table 2a), and 2) the subset of mice that had grooming, open field, rotarod, and sociability data (“subset of mice that have sociability data”)(see Table 2b).

**Table 2:** Tables depicting the sample size in each strain, broken down by treatment. A minimum of n=14 was aimed for per genotype, per strain, per treatment. Table (a) shows the samples for all behaviors except sociability. Table (b) shows the reduced sample size due to failed acquisition in the sociability tri-chamber test.

| (a) Totals- Behavior |  |  |  | (b) Totals- Sociability |  |  |  |
| --- | --- | --- | --- | --- | --- | --- | --- |
| Strain | Total | Oxytocin | Placebo | Strain | Total | Oxytocin | Placebo |
| *16p* | 30 | 15 | 15 | *16p* | 14 | 9 | 5 |
| 129 | 29 | 15 | 14 | 129 | 17 | 10 | 7 |
| *Fmr1* | 29 | 14 | 15 | *Fmr1* | 18 | 8 | 10 |
| FVB | 30 | 13 | 16 | FVB | 18 | 10 | 8 |
| *Shank3* | 36 | 17 | 19 | *Shank3* | 29 | 13 | 16 |
| C57 | 26 | 12 | 14 | C57 | 19 | 10 | 9 |

#### 3.5.2 Behavioral Tests

A description of each of the behavioral tests follows (Crawley (2007); Silverman et al. (2010)), in the order they were performed:

##### Sociability

The three-chambered sociability apparatus consists of a rectangular box with three chambers, with a wire cup placed in either side chamber. The mouse is habituated for 10 minutes during which it can access all three chambers. After this time, a novel stranger mouse is placed under a wire-cup in one of the side chambers. The mouse is observed for an additional 10 minutes during which measures such as time (s) spent in each of the three chambers, specifically near/sniffing the wire cups (near the novel mouse) are observed using an overhead video system, manually scored, and then compared. Mice that are not considered social spend either an equivalent amount of time between the two cups or show a preference for the empty cup (away from the novel mouse).

##### Rotarod

The rotarod test was run on a system (Rota-rod/RS, Panlab, Harvard Apparatus) with the capacity to run up to five mice at a time (though no more than three were ever run). For the rotarod test, mice go through two types of trials, pretrials and test-trials, over two days. Pre-trials are sessions where the mice are allowed to learn how to stay on the rod for 10s, when it is only rotating at 4 revolutions per minute (rpm). The number of attempts it takes before the mouse stays on the rod for the entire duration is recorded. Latency to fall off the rod is the primary measure of the rotarod test. The mice are given three trials of a total possible five minutes for each trial. The rod starts off at 4rpm and slowly accelerates over the course of 5 minutes. The latency (s) to fall off the rod is recorded, averaged, and compared. Mice that either have a learning and memory deficit or a motor deficit, perform worse on one or both measures when compared to wildtype controls.

##### Grooming

For the grooming test, the mice are placed into an empty standard cage (Green Line IVC Sealsafe PLUS Mouse, Techniplast) for ten minutes (habituation). The following ten minutes, mice are placed into a different, empty, standard cage and observed for grooming behaviors. Primary measures are duration of grooming bouts (s) as well as number of grooming bouts. Mice with a repetitive behavior phenotype spend significantly more time grooming, with more bouts of grooming, than wildtype controls.

##### Open Field

The open field test consists of 10 minutes of testing, where the mouse is placed in an empty box (area=44cm^2^) and allowed to roam free for the duration of the test. The mouse’s motion is recorded by a system (Activity Monitor 7, Med Associates Inc, Fairfax VT) and primary measures investigated are time (s) spent in the center (decreased anxiety) and total distance (cm) moved (hyperactivity).

It should be noted that mice underwent a fifth behavioral test during the study (after the other behavioral tests), the novel object recognition test. This data was removed from the analysis in it’s entirety once it was discovered that the tracking software was not accurately capturing the mouse’s behavior.

## 4 Results

### 4.1 Chronic intranasal oxytocin treatment has a trending effect on the neuroanatomy of autism-related mouse models

Chronic treatment with oxytocin had a weak and focused effect, only changing the volume of the cerebellar peduncle superior and the endopiriform claustrum (intermediate nucleus) regions when assessing relative volumes (see Figures 3 and 4), across all mouselines (using Equation 2). No other brain regions were significant with correction for multiple comparisons, but the pattern of treatment effect, as shown by the voxel-wise analysis, shows alterations in several cerebellar regions, the cingulate cortex, and, interestingly, several regions related to sensory/motor behaviors, like the cuneate nucleus, the dorsal tenia tecta, and the superior olivary complex. Interestingly, OXTR gene expression did not correlate with neuroanatomical changes due to oxytocin treatment (based off Equation 1).

**Figure 2:**
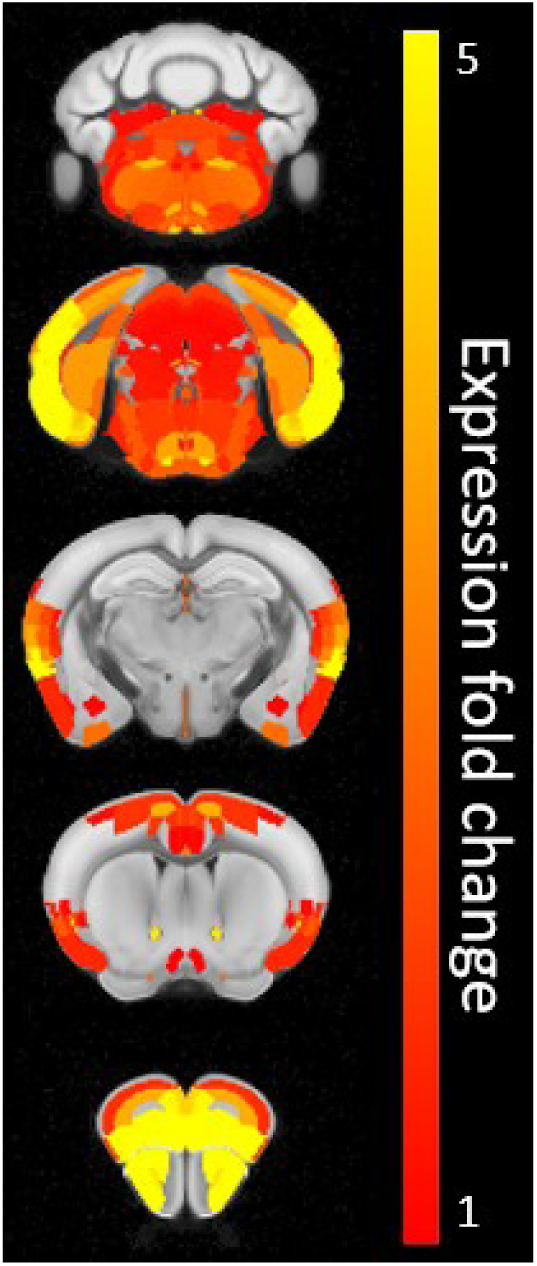
Figure depicting OXTR gene expression information in the adult mouse brain overlaid on a neuroanatomical map to depict how OXTR gene expression energy varies (yellow is greatest). We would expect the greatest volume changes from oxytocin treatment to occur in areas with the highest oxytocin affinity. In the brain, OXTRs are most abundant in the olfactory system, the central nucleus of the amygdala, the CA1 region of the hippocampus, the ventromedial hypothalamus, and the nucleus accumbens (Benarroch (2013); Gould and Zingg (2003)). Gene expression data was gathered from the Allen Institute, following methods previously described (Carriere et al. (2020); Fernandes et al. (2017); Lein et al. (2007); Lindenmaier et al. (2021); Sunkin et al. (2012); Yee (2019))

**Figure 3:**
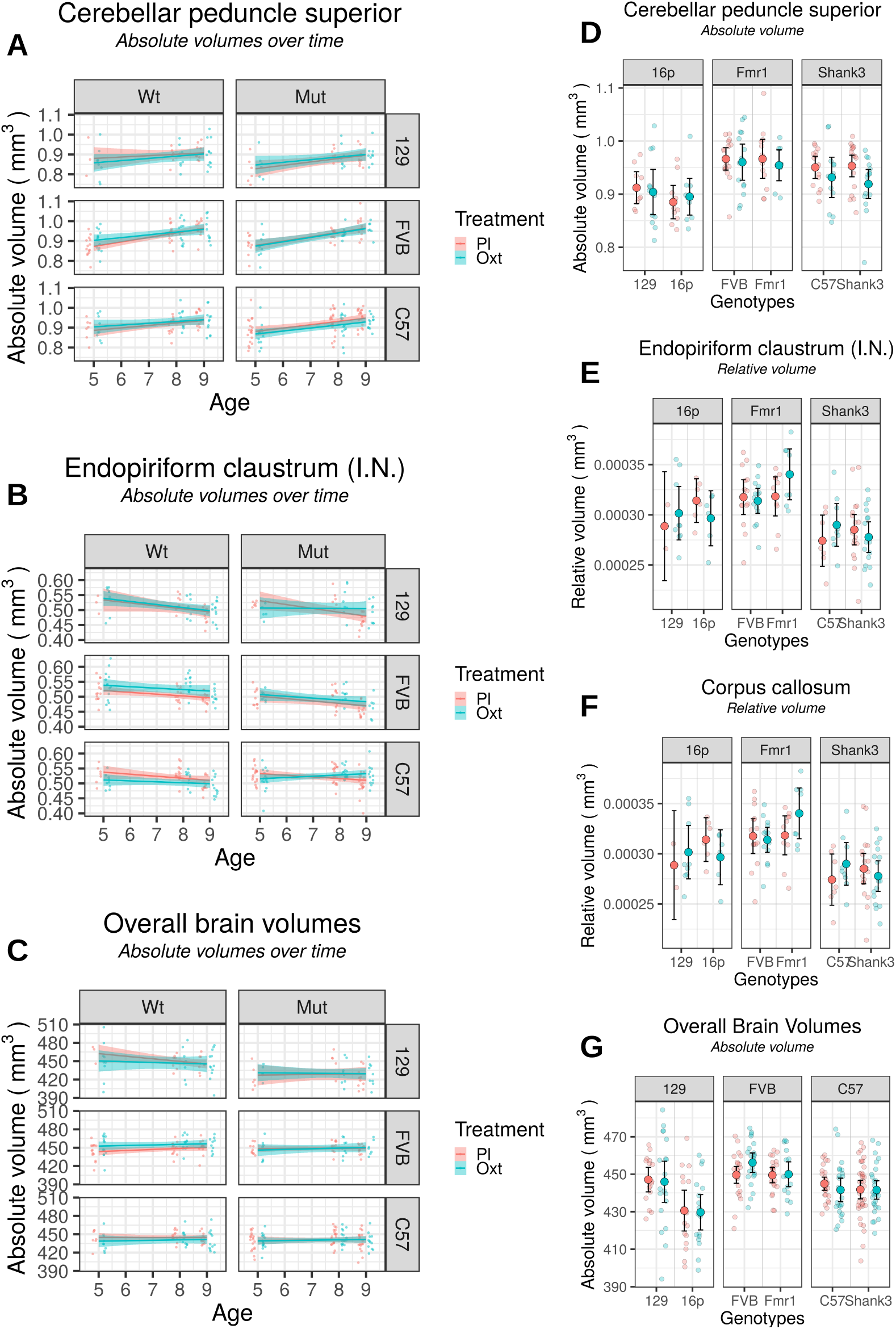
Figures depicting treatment effects of selected regions from *in vivo* imaging. Figures **A** through **C** show the change in volume over time of the cerebellar peduncle superior, the endopiriform claustrum, and the total brain volumes. Figures **D** through **G** show a cross-sectional view of the brain region volumes after 3-4 weeks of treatment, faceted by strain. To better see the treatment effect, two of the figures (**E** and **F**) show relative volume of the regions. With correction for multiple comparisons, no significant treatment effect was seen on any region, though the cerebellar peduncle superior and endopiriform claustrum (intermediate nucleus) showed trending (q=0.155) effects (see 4). There were significant genotype differences across brain regions, and overall brain volume.

**Figure 4:**
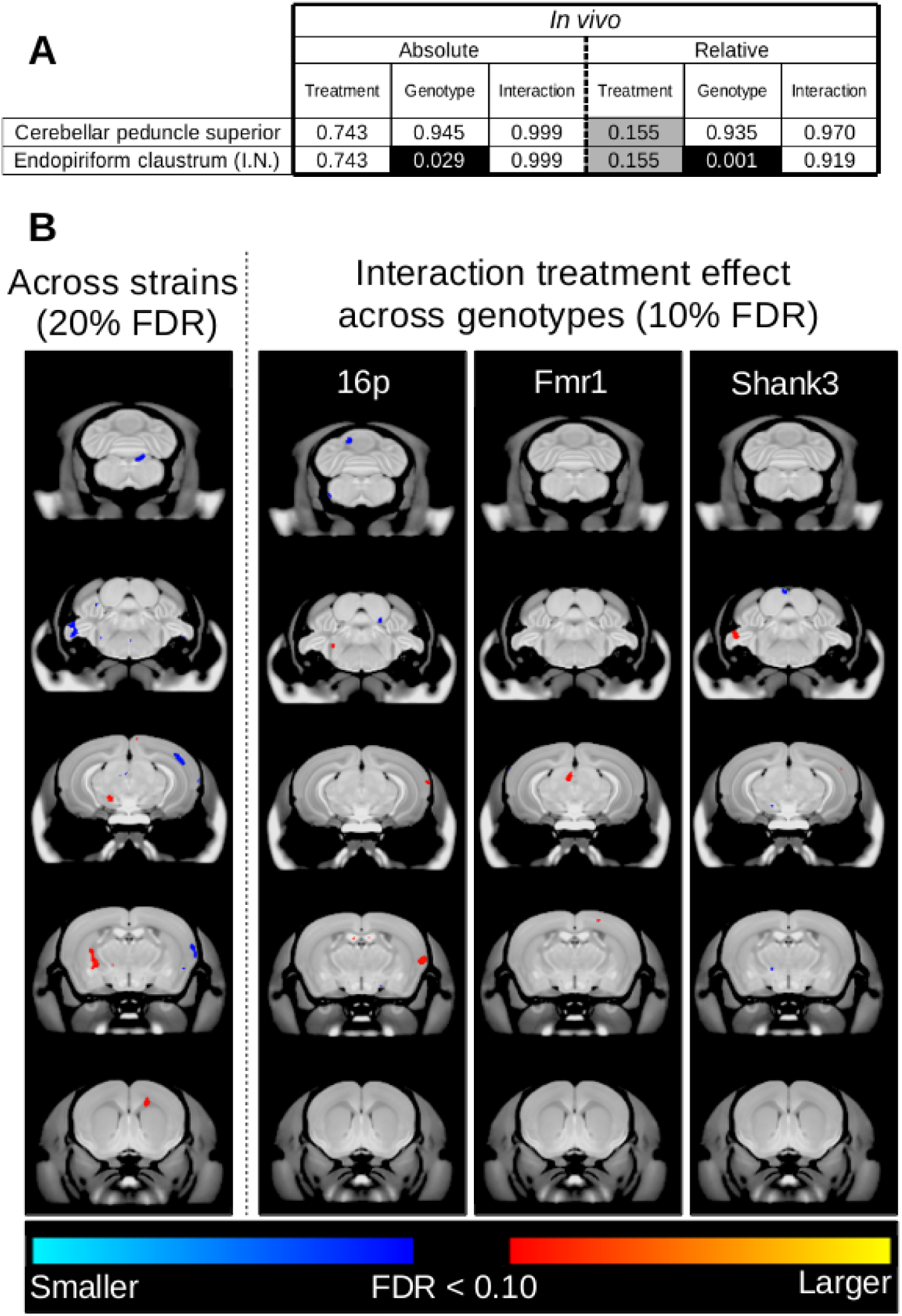
Treatment effect in the *in vivo* data. Top (**a**) shows the results of the volume-wise linear mixed effects model with main effects of treatment and genotype, with a random intercept for each background strain (using Equation 2). FDR-corrected values are shown with black background indicating significance (q <10%) and grey background showing trends. Results from both absolute and relative volume analyses are shown; in this case, relative volumes of each region were calculated as proportion of the total brain volume. The cerebellar peduncle superior and the endopiriform claustrum (I.N. = intermediate nucleus) showed subtle treatment effects. Bottom (**b**) shows the voxel-wise data analysis, using the linear mixed effects model shown in Equation 1. Colors indicate voxels that were q <10% significant, with red indicating regions that were larger than the reference group and blue indicating smaller. In the case of the treatment effect across strains (left), colours indicate q <20%, to better distinguish patterns. These figures clearly depict the subtle effect of treatment response on neuroanatomy.

The *ex vivo* analysis (using Equation 1), showed no significant treatment effect (see Figure 8), only a weak trend (q = 0.195) in the *Shank3* interaction of cerebellar lobule 9 (white matter). When assessing strain differences (using Equation 3), the *Shank3* mice showed trending (q = 0.161) treatment effect in three regions of the primary somatosensory cortex and the lateral ventricles. As seen in Figure 8, the genotype differences were significant and widespread, although more so for the *16p* and *Fmr1* mice, with significant effects in regions such as the midbrain, hippocampus, and cerebellum in the *16p* mice, in regions including the hippocampus, cerebellum and white matter tracts in the *Fmr1* mice, and in the cerebellar peduncle, habenular commissure, and pontine nucleus for the *Shank3* mice.

### 4.2 Chronic intranasal oxytocin treatment has little to no effect on the behavioral phenotype of autism-related mouse models

No significant effect of oxytocin treatment was found on social behavior, regardless of strain or genotype when analyzing the sample as a whole (Figure 6A and E). A significant effect of treatment was found when subsetting for the FVB background strain, with treatment normalizing a deficit in the number of bouts of grooming in the *Fmr1* mouse model (q=0.012)(Figure 6B). No other effects survived correction for multiple comparisons (see Figure 7A).

**Figure 5:**
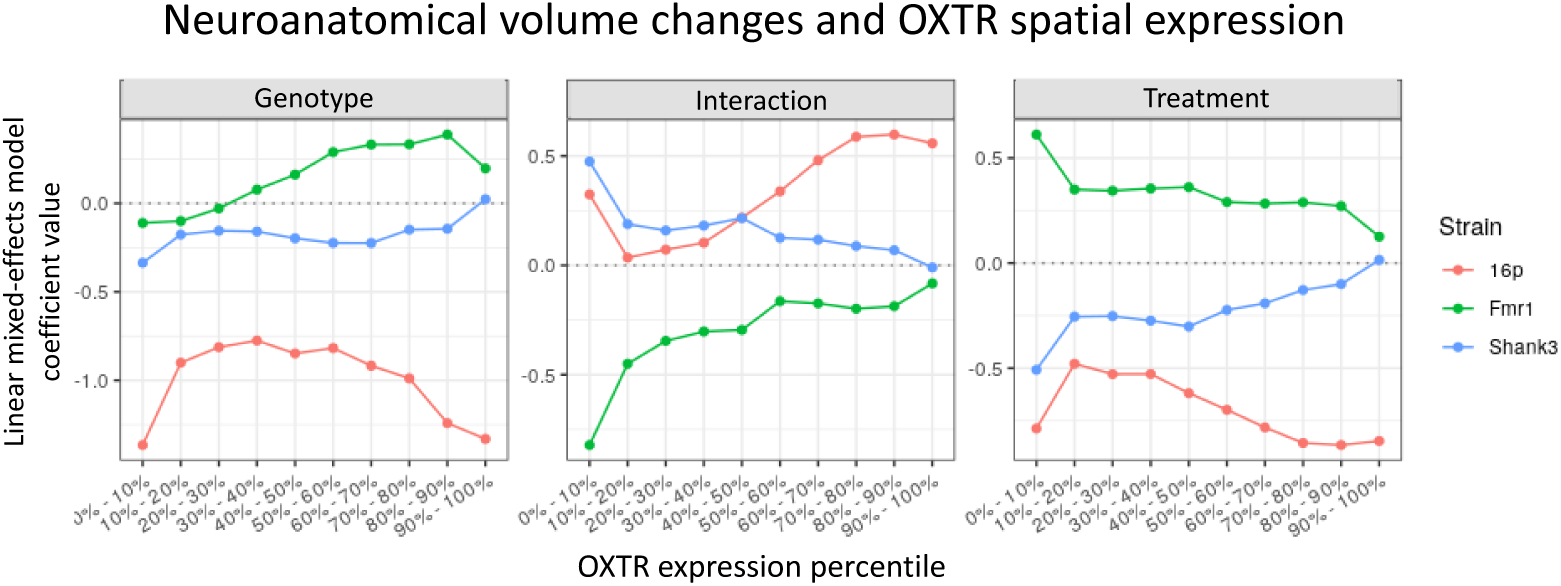
Figure depicting neuroanatomical volume changes and OXTR spatial expression. Specifically, the brain was divided into 10 regions based on spatial OXTR expression patterns in the adult mouse brain. The coefficient value from the linear mixed-effects model from Equation 3 for each subdivision is shown, for each mouse model (colors), by each variable of interest (facets). The coefficient represents how much the brain changes in (absolute) volume due to the variable of interest (i.e. Treatment (right)), in the specific region of interest. For example, in the case of the Fmr1 and Shank3 mouse models, with increasing OXTR expression, the volume change due to treatment approaches zero, opposite to expectation.

**Figure 6:**
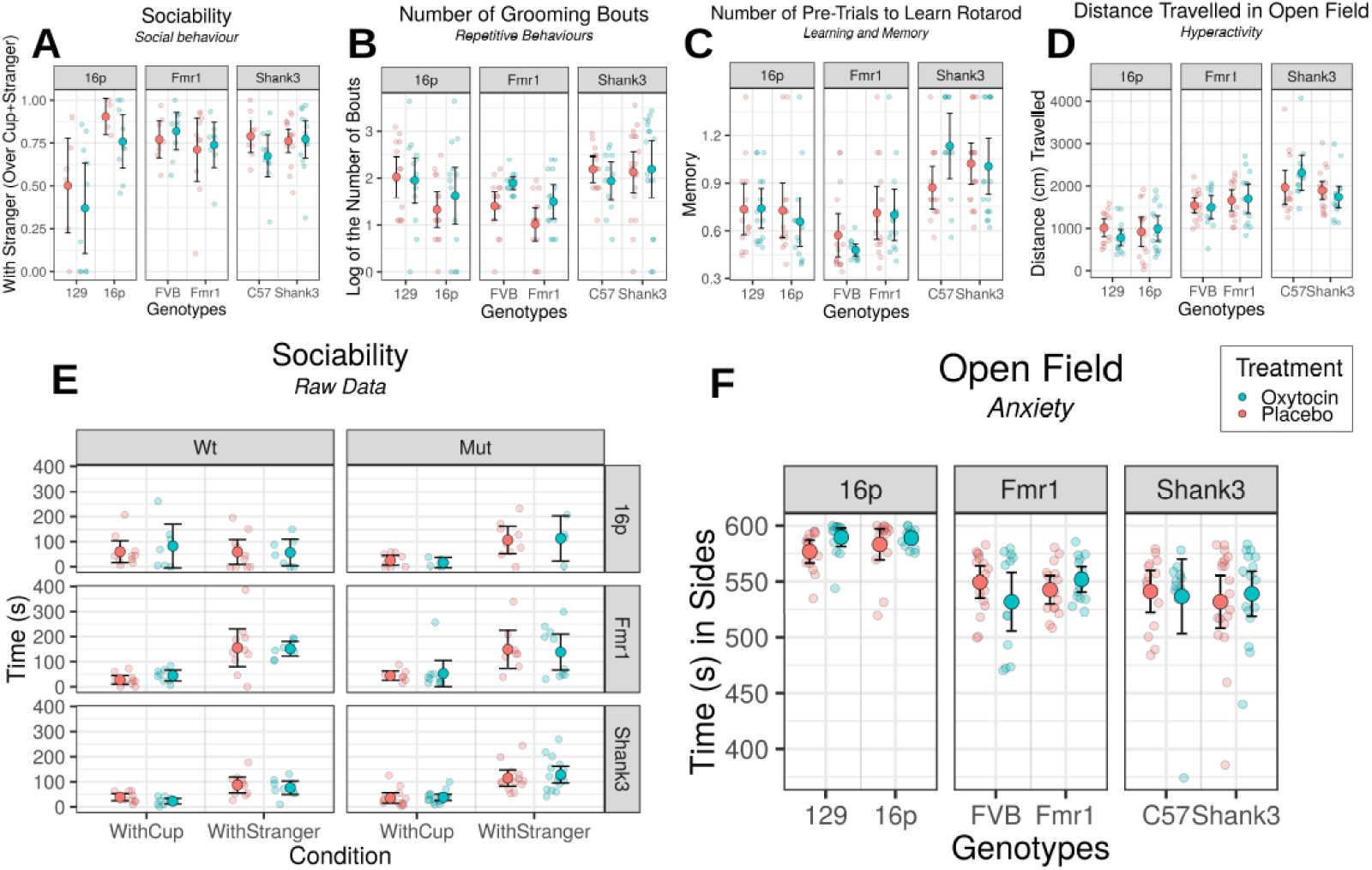
Figures depicting treatment effect in each strain, by genotype. Blue indicates mice treated with oxytocin, red indicates mice treated with saline (placebo). Each point represents one mouse. Bars show mean ± confidence interval (95%). Sociability data is shown both as (**E**) the raw time spent with the mouse and cup, for each mouse by genotype and strain, as well as (**A**) a ratio of time with stranger over total time with cup and stranger. (**B**) Number of bouts of grooming in the mice, (**C**) number of pre-trials required to learn rotarod, (**D**) distance travelled in the open field, and (**F**) time spent in the sides (= *total − center*) of the open field are also shown.

**Figure 7:**
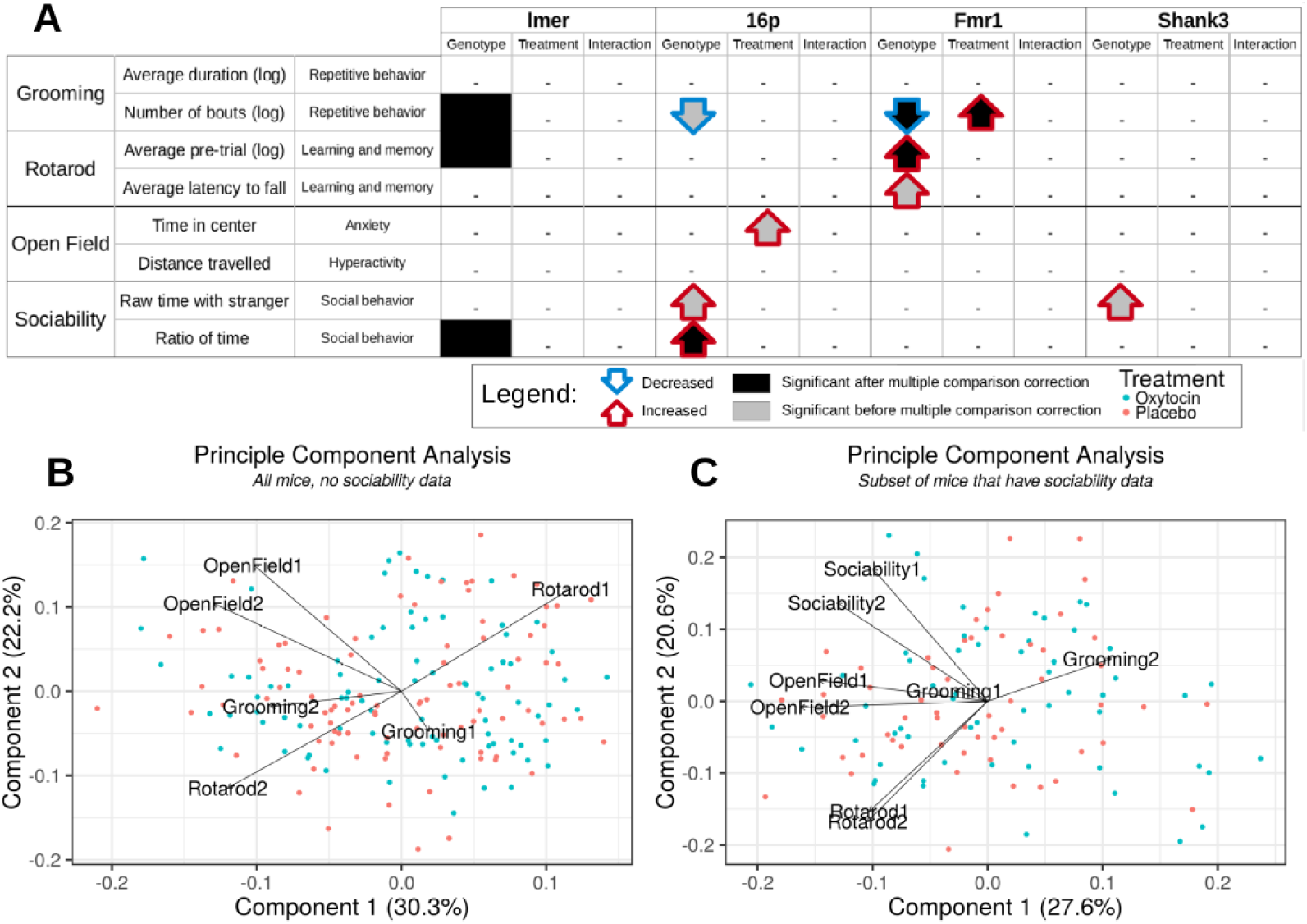
Table **A** shows the results of the log-likelihood tests assessing the main effects of treatment and genotype, and their interaction from the linear mixed effects models on the behavioral tests. Each of the behavioral tests are shown, broken down by measure. The table is color-coded to show significant, corrected q values (<0.05) with a dark background, and trending (q <0.10) values with a grey background. Arrows indicate changes relative to their respective controls (i.e. *16p* mice to 129 mice). Figures **B** and **C** show the results of the PCA analyses, with the “all mice, no sociability data” set on the left (**B**) and the “subset of mice that have sociability data” on the right (**C**). The names of the PCA measures follow down along the table of measures. Specifically, Grooming1 is “average duration” (the first row in the table), Grooming2 is “number of bouts” (second row), Rotarod1 is “average pre-trials” (3rd row), Rotarod2 is “average latency to fall” (4th), OpenField1 is “time in center” (5th), OpenField2 is “distance travelled” (6th), Sociability1 is “raw time with stranger” (7th), and Sociability2 is “ratio of time” (last row). As evident from these tables and figures, little to no treatment effect was observed across the multitude of behavioral phenotypic measures employed in this study.

**Figure 8:**
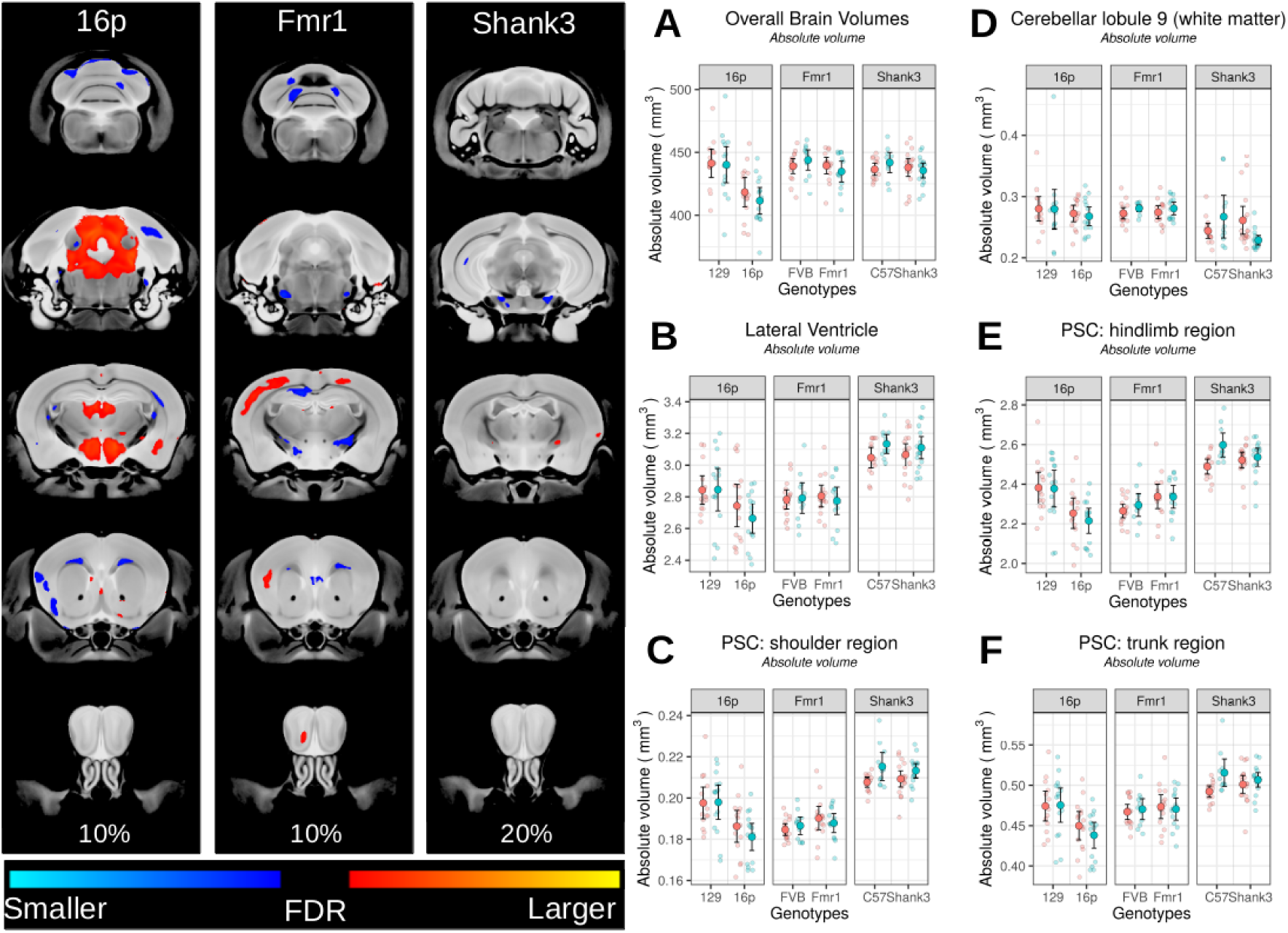
Figures depicting the results of the *ex vivo* analyses. On the left, genotype effects are shown from the subsetted (by background strain) voxelwise analyses (Equation 3). On the right, the graphs depict treatment effect across strains for six regions, including overall brain volumes. PSC= Primary Somatosensory Cortex

Without correction for multiple comparisons, when subsetting for background strain, some trends for treatment were observed. Treatment appeared to decrease the amount of time spent in the center of the open field in the *16p* (p=0.048) and the *Fmr1* (p=0.086, non-significant interaction) mice (Figure 6F). Treatment also had an interesting trending effect on the C57 mice (*Shank3* control), by decreasing performance on the rotarod pretrials (interaction p=0.070)(Figure 6C) and increasing hyperactivity in the open field (interaction p=0.093), while not having an effect on the *Shank3* mutant mice (Figure 6D)

PCA was employed to further elucidate if a treatment effect was present that was not captured in the individual behaviors but would be captured given a multivariate analysis. Two analyses were run on separate subsets, the first on the full data-set excluding social behaviors (“all mice, no sociability data”)(see Table 2a), and the second on the subset of mice that had social behaviors tested (“subset of mice that have sociability data”)(see Table 2b). The first two components accounted for 52.5% and 48.2% of the total variance, respectively, of the two datasets (see Figure 7B,C). Most of the behavioral tests exhibited negative values for principle component 1 (PC1), in both sets. In contrast, the pre-trials measure of rotarod and the average duration of grooming bouts were positive in both sets. No distinct groups of variables were distinguishable, although measures from the same behavioral test did coincide. Overall, no pattern of treatment response was observed.

## 5 Discussion and Conclusions

Chronic treatment with intranasal oxytocin produced subtle behavioral and neuroanatomical effects across three different autism mouse models. Treatment effect on the neuroanatomy was modest, and primarily apparent when assessing relative volumes. No significant effect of treatment was found on social behavior, although a significant effect of treatment was found in the *Fmr1* mouse, with treatment normalizing a grooming deficit. No other treatment effect on behavior was observed that survived multiple comparisons correction. No pattern of treatment effect was distinguishable when a multivariate method (PCA) was employed.

Although not statistically significant, the pattern of treatment effect in the brain involved many regions that are associated with oxytocin signaling and, moreover, have been implicated in autism pathogenesis. For example, the brain stem and olfactory systems have large populations of oxytocin receptors, and we found (trending) treatment effects in the cuneate nucleus, the superior olivary complex, and the dorsal tenia tecta, among other regions. However, contrary to expectation, OXTR expression did not correlate with neuroanatomical changes due to oxytocin treatment.

The superior cerebellar peduncle– a region found to have a trending treatment effect– is a white matter tract that connects the cerebellum to the midbrain. A number of studies have shown aberrations in the structure and function (see Crippa et al. (2016) for review, also Cheng et al. (2010)) of the superior cerebellar peduncle in autism. Moreover, the cerebellum, as a whole, has been implicated in the ethiopathogenesis of autism (for review, see Fatemi et al. (2012), Crippa et al. (2016)). An elegant study in a mouse model of autism demonstrated the important role of the cerebellum in autism-related behaviors, including the therapeutic potential of cerebellar neuromodulation in autism (Tsai et al. (2012)).

The endopiriform claustrum, another region that was found to have a trending treatment effect, is a part of the basal ganglia, the system known to regulate repetitive behaviors. Although the exact function of the claustrum is unknown, it is hypothesized that it plays a role in multisensory integration and is an important relay nucleus to the cortex (Mathur (2014)). Furthermore, a number of regions in the *Shank3* mouse model found to be affected by treatment, are related to sensation/perception. There is a wealth of literature on the dysfunction of sensation/perception in autism (see Robertson and Baron-Cohen (2017) for review). It’s important to acknowledge that these effects were all modest and more study is necessary to truly assess the potential promise of instranasal oxytocin in treating autism.

To compare to existing literature, the effect of genotype of the three mutant mice was separately analyzed (see Equations 5 and 3). Without correction for multiple comparisons, all three mutant mice, for the most part, showed behavioral and neuroanatomical differences that were expected based off previous literature (Bernardet and Crusio (2006); Bozdagi et al. (2010); Ellegood et al. (2015); Harony-Nicolas et al. (2017); Horev et al. (2011); Yang et al. (2012)). With multiple comparison correction, this study moderately recapitulated existing literature. Additional analysis comparing to data distributions from previous literature and previous studies within our lab was conducted after the primary results were analyzed, to demonstrate the lack of effect of methodology on the behavior of the mice.

Possible explanations for the lack of treatment effect, particularly in the behavioural measures, include a translation hurdle from acute to chronic oxytocin treatment, study population differences, and dosage.

Some studies investigating the translational hurdle from acute to chronic treatment have found value in the glutamatergic system, the genetic background of the oxytocin receptor, and in the dosage. A recent study comparing acute and chronic treatment in both mice and humans suggests a unique sensitivity of the glutamatergic system to repeated oxytocin administration may explain the differential behavioral effects of oxytocin between the two lengths of administration (Benner et al. (2021)). Another study that did not find improvements with oxytocin effect in the larger study population, did find significant effects in a subgroup of participants. Their results suggest that efficacy of long-term oxytocin administration in young men with high-functioning autism depends on the oxytocin dosage and the genetic background of the oxytocin receptor (Kosaka et al. (2016)). This further exemplifies the role neuroimaging could play in distinguishing subtypes of autism.

Another possible issue for the lack of treatment effect may be due to the study population. We chose to limit the study to older (adolescent/adult) mice to minimize variability and increase our power. However, it is possible that a younger population is needed to see the effects of oxytocin. For example, the association of the plasma oxytocin levels in autism (that was mentioned earlier as a promising reason we expect response to oxytocin) was found in children, and perhaps does not extend to adults (Modahl et al. (1998)). Reversal of some abnormalities have been observed in Shank3 deficient mice in early stages of development with (acute) oxytocin treatment (Reichova et al. (2020)). Similarly, one of the only clinical trials that saw improvements with chronic oxytocin treatment was conducted in children (Yatawara et al. (2016)). Developmental processes occur during childhood that could impact the effect of treatment response (Bhardwaj et al. (2006); Pascual-Leone et al. (2011); Sorrells et al. (2018)).

Along these lines, the three mouse lines employed were primarily chosen for their high face validity to autism, and not necessarily for their utility for treatment with oxytocin. Specifically, without an underlying deficit in a particular behavior, no effect of oxytocin on behaviour is expected, regardless of promise. The lack of social deficits in all three mouse lines found in this study obscures the possible treatment effect of oxytocin on social behavior. In contrast, neuroanatomical imaging has greater sensitivity and translational potential for understanding circuit changes.

Moreover, some other limitations arose during this study. For example, the timeline was designed to maximize throughput and therefore some potentially insightful behavioral tests were omitted. Given the number of brain regions involved in sensation/perception that showed changes due to treatment, a test that specifically evaluated those behaviors may have seen a significant effect of treatment. The reduced sample size (Table 2) for the sociability test (due to failed acquisition) could have also contributed to the lack of treatment effect, due to low power.

Furthermore, there is no current robust method to assess the quantity of oxytocin that is delivered to the brain using the intranasal method. When the treatment was first started, all the mice fought the scruff hold and intranasal administration equally. Near the end of the treatment period, however, approximately half the mice would fight the treatment while the other half remained stationary. The authors assume the mice that ceased avoiding the intranasal application were the ones to receive oxytocin and, through classical conditioning, had learned that the application would quickly lead to pleasant sensations (Benarroch (2013)). Though this was primarily observational, a small sample of mice (n=14) were assessed to see if treatment arm (placebo vs. oxytocin) could be determined just through interaction during treatment after 3 weeks of treatment. With the help of a colleague, the correct treatment arm was confirmed for all the mice in the sample. Although not specific to the brain, this confirms that the intranasal application method was successful in administering oxytocin into the mice.

Along these lines, it is important to consider the dosage employed in this study. Although chosen to be relevant to the clinic, it is on the upper end of doses used in previous studies (Bales et al. (2013); Guastella and Hickie (2016); Neumann et al. (2013); Quintana et al. (2019)). Recent literature suggests an inverted U-curve doseresponse relationship of oxytocin, and therefore the promise of oxytocin treatment can not be dismissed given the single dose investigated in this study (Owada et al. (2019)). Similarly, previous studies suggest a potential deterioration of efficacy of chronic treatment, indicating that perhaps the treatment period in this study was too long to detect effects of oxytocin (Benner et al. (2021); Owada et al. (2019)). Yet another study found a time-course change of efficacy of oxytocin on social behaviors in adults with autism (Owada et al. (2019)), indicating the dose used in this study may be too low or too high, for too long of a period of time. Further studies on these factors may lead to a better understanding of the best protocol to see a significant treatment effect of oxytocin. As discussed previously, it remains possible that the treatment effects observed were solely due to peripheral treatment effects (Leng and Ludwig (2016)). More tests that assess the utility of intranasal oxytocin, like proper dose-response studies, with peripheral effects controlled for, are necessary (Galbusera et al. (2017); Leng and Ludwig (2016); Pagani et al. (2020)).

Future studies should aim to explore other behavioral tests, population sets, and administration methods. The authors recognize that the exclusion of female mice from this study contributes to the ongoing sex bias in neuroscience and this study should be replicated in female mice to truly assess the utility of oxytocin administration in autism mouse models.

The possibility remains that the three autism mouse models employed in this study– the *16p, Fmr1*, and *Shank3* mouse models– are non-responders to oxytocin. Despite the limitations, the study is one of few studies to assess the effects of chronic oxytocin treatment on the neuroanatomy and behavior of multiple mouse models of autism.

## 6 Declarations

### 6.1 Ethics approval and consent to participate

All studies and procedures were approved by The Centre for Phenogenomics (TCP) Animal Care Committee, in accordance with recommendations of the Canadian Council on Animal Care, the requirements under the Animals for Research Act, RSO1980, and the TCP Committee Policies and Guidelines.

### 6.2 Availability of data and material

The datasets from this study are available from the corresponding author on reasonable request.

### 6.5 Authors’ contributions

Conception and interpretation of this project were done by ZL, JE, EA, and JPL. Design was primarily ZL, JE, JF, EA, and JPL, while acquistion/analysis was primarily ZL, MS, KE. Additionally, ZL, JE, KE, and JPL contributed to the writing or revision of the manuscript, and all authors read and approved the final manuscript

## 7 Acknowledgements

The authors would like to thank Darren Fernandes, Chris Hammill, Dr. Dulcie Vousden, and Matthijs Van Eede for helping with data analysis, Christine Laliliberte, Dr. Dulcie Vousden, and Dr. Jill Silverman for procedural assistance, and Drs. Brad Wouters and Mark Henkelman for advice and guidance.

